# Ultrasensitive Dopamine Detection with Graphene Aptasensor Multitransistor Arrays

**DOI:** 10.1101/2022.03.04.483061

**Authors:** Mafalda Abrantes, Diana Rodrigues, Telma Domingues, Siva S. Nemala, Patricia Monteiro, Jérôme Borme, Pedro Alpuim, Luis Jacinto

## Abstract

Dopamine is a neurotransmitter with critical roles in the human brain and body, and abnormal dopamine levels underlie brain disorders such as Parkinson’s Disease, Alzheimer’s Disease, and substance addiction. Herein, we present a novel high-throughput biosensor based on graphene multitransistor arrays (gMTAs) functionalized with a selective aptamer for robust ultrasensitive dopamine detection. The miniaturized biosensor integrates multiple electrolyte-gated graphene field-effect transistors (EG-gFETs) in an array configuration, fabricated by high-yield reproducible and scalable methodologies optimized at the wafer level. With these gMTA aptasensors, we reliably detected ultra-low dopamine concentrations in physiological buffers, including undiluted phosphate-buffered saline (PBS), artificial cerebral spinal fluid (aCSF), and high ionic strength complex biological samples. We report a record limit-of-detection (LOD) of 1 aM (10^−18^) for dopamine in both PBS, and dopamine-depleted brain homogenate samples spiked with dopamine. The gMTAs also display wide sensing ranges across all media, up to 100 μM (10^−8^), with a 22 mV/decade peak sensitivity in aCSF. Furthermore, we show that the gMTAs can detect minimal changes in dopamine concentrations in small working volume biological CSF samples obtained from a mouse model of Parkinson’s Disease.

## 1. Introduction

Neurotransmitters are molecules indispensable for the communication between neurons and play a critical role in brain function. The accurate detection of neurotransmitters concentrations in the brain and biological samples is of great importance for neurobiology research and developing novel diagnostic and therapeutic approaches for brain disorders affecting neurotransmitter levels and dynamics. A significant neurotransmitter is dopamine (3,4-dihydroxyphenethylamine), which has essential roles in the human brain and body, regulating several physiological processes involved in motor function, memory, motivation, arousal, and reward [1]. Abnormal alterations in the levels of dopamine can have severe consequences and underly brain disorders such as Parkinson’s Disease (PD), Alzheimer’s Disease, Schizophrenia, Attention Deficit and Hyperactive Disorder, and substance addiction [1]–[3]. The ability to detect physiologically relevant dopamine concentrations by high-throughput approaches in the brain or brain-derived biological samples can accelerate the development of early diagnostics and improved therapeutics for these disorders. However, conventional analytical methodologies to monitor and detect dopamine, which include enzyme-linked immunosorbent assays (ELISA), high-performance liquid chromatography (HPLC), capillary electrophoresis, and spectroscopy, rely on large-scale, expensive equipment, or require laborious sample preparation and long detection cycles [4], [5]. Novel emerging approaches have focused on miniaturized biosensors, primarily based on catalytic and electrochemical reactions, but have mostly lacked relevant selectivity and sensitivity [6], [7]. At present, the limit of detection (LOD) for dopamine sits at 0.5 fM [8], [9], but most methods have reduced sensitivity, and narrow working ranges between nM and μM concentrations [6], [7]. Additionally, many novel biosensors for dopamine and other neurotransmitters detection have complicated fabrication processes [6], [7] that pose constraints on miniaturization and integration, thus limiting dissemination, reproducibility, and the development of relevant research tools and point-of-care devices.

Graphene-based biosensors have been attracting growing attention due to their extremely high sensitivity based on graphene’s unique electronic properties, high chemical and mechanical stability, and biocompatibility [10]–[13]. Graphene field-effect transistors (gFETs), in particular, take advantage of graphene’s exceptionally high carrier mobility and surface-to-volume ratio to permit high signal-to-noise transduction of biodetection events through electrostatic gating [13]–[15]. Because the gFETs’ transduction depends on the field-effect modulation based on different local doping mechanisms, charge carrier scattering, and dielectric environment [16]–[18], they can be designed and tuned according to application demands. The graphene channel on gFETS can also be functionalized through surface chemistry with biorecognition elements such as enzymes, antibodies, DNA, and aptamers for selective biodetection [10], [19]–[21]. A wide range of selective biosensing applications, including protein and DNA detection in optimized buffers and samples prepared from body fluids such as blood, sweat, or saliva, has been reported in the last 10 years [11], [22], [23], including by us [22], [24]. However, despite promising operation in controlled conditions, gFET biosensors, like any ion sensitive FET, suffer from dramatic sensitivity reductions in biological conditions or media. The decreased sensitivity occurs due to the reduction of the Debye length [25], i.e., the distance over which the local electric field can modulate charge carriers in the graphene channel, due to ionic strength increase. Approaches to overcome these limitations in FET-based biosensors have included electronic tunning by adding a floating gate configuration [26], removal of the surface excess ion population [27], channel morphology alterations [28], [29], and reduction of the biorecognition element size [28], with the latter being a promising approach for gFET biosensors to preserve crystalline graphene’s unique properties without considerably increasing fabrication complexity.

Additionally, many of the recently proposed graphene biosensors do not address issues relating to replicability and scalability of fabrication methods and the stability and reproducibility of the desired measurements. Thus, the deployment of graphene-based point-of-care devices and reliable academic and pharmaceutical research tools has been limited despite significant efforts. Wafer-scale graphene synthesis and fabrication of graphene biosensors with replicable methodologies can improve device uniformity and facilitate integrated designs with multiplexed parallel-assays for high-throughput measurements [10], [30]–[33], a desired feature for real-world neurotransmitter sensors.

This work proposes a novel graphene biosensor that permits ultrasensitive and selective dopamine detection, yielding stable and reliable results by high-throughput sample measurement replication. Micron-sized electrolyte-gated field-effect transistors (EG-gFETs) were fabricated with reproducible methodologies optimized at wafer level and integrated into a miniaturized multi transistor array (gMTA). This array was converted into an aptasensor by functionalizing EG-gFETs with a dopamine-specific aptamer, promoting the redistribution of charges within the Debye length. Taking advantage of multiple parallel sample measurements in a single gMTA, the high mobility of crystalline graphene’s charge carriers, and effective screening of the aptamer’s charge redistribution upon dopamine binding, we report a record LOD for dopamine with a wide sensing range. Furthermore, we show that these gMTAs can detect minimal changes in dopamine concentration in small working volume biological cerebrospinal fluid (CSF) and brain homogenate samples from an animal model of PD. This feature is highly pertinent for developing novel point-of-care devices and research tools that require stable high-throughput detection of physiologically and clinically relevant dopamine concentrations.

## 2. Results and Discussion

### 2.1. Fabrication of graphene multi-transistor arrays (gMTA)

High-throughput ultrasensitive detection of dopamine in real-world biosensing applications requires the realization of a miniaturized biosensor that can yield multiplexed stable measurements in a wide range of sensing media and contexts. Reproducible wafer-scale high-yield fabrication methods are also required to ensure the replicability of graphene-based biosensors. Therefore, an Si/SiO_2_ wafer containing 784 graphene multi transistor arrays (gMTAs) with 15680 electrolyte-gated graphene field-effect transistors (EG-gFETs), with a yield of approximately 80%, was fabricated for this work (Fig. 1A). Each 4.5 × 4.5 mm^2^ gMTA chip consists of an array of 20 EG-gFETs with individual gold drain electrodes connected to 2 common gold source electrodes, with groups of 10 transistors sharing a common source, and respective interconnect lines (Fig. 1B,C). This design leverages our previously published architecture, including an integrated co-planar electrolytic gold gate electrode [34] that removes the necessity of a cumbersome external floating gate electrode for modulation of the local electric field. This integration facilitates the fabrication of compact and portable devices for real-world applications. Drain and source electrodes are connected by a 25 μm long and 81 μm wide single-layer graphene channel (Fig. 1C). Graphene was grown by chemical vapor deposition (CVD), commonly used to produce large-area polycrystalline single-layer graphene sheets [35] (Supporting Informatio, Fig. S1), and patterned by optical lithography. Dielectric passivation of source and drain electrodes was achieved with a 250 nm multi-stack passivation layer of SiO_2_/SiN_x_ that improves the impermeability to solvents during the following stages of transistors’ functionalization, increasing resistance to delamination in prolonged exposure to liquid solutions for biosensing applications [36]. Finally, the gMTA chips were mounted and wire-bonded to a custom-designed PBC for electronics interfacing (Fig. 1D). The overall fabrication method is highly reproducible at a low cost and optimized at the wafer level to preserve graphene’s electronic properties. The integrated multitransistor array design allows simultaneous parallel measurements in different EG-gFETs in one single chip. This parallel multi-sampling approach provides redundant measurements that produce robust averaged data from single samples. The variance in the EG-gFETs response in each gMTA is also measured, and mal-functioning transistors can be disconnected without contributing to the final readout values.

**Figure 1.**
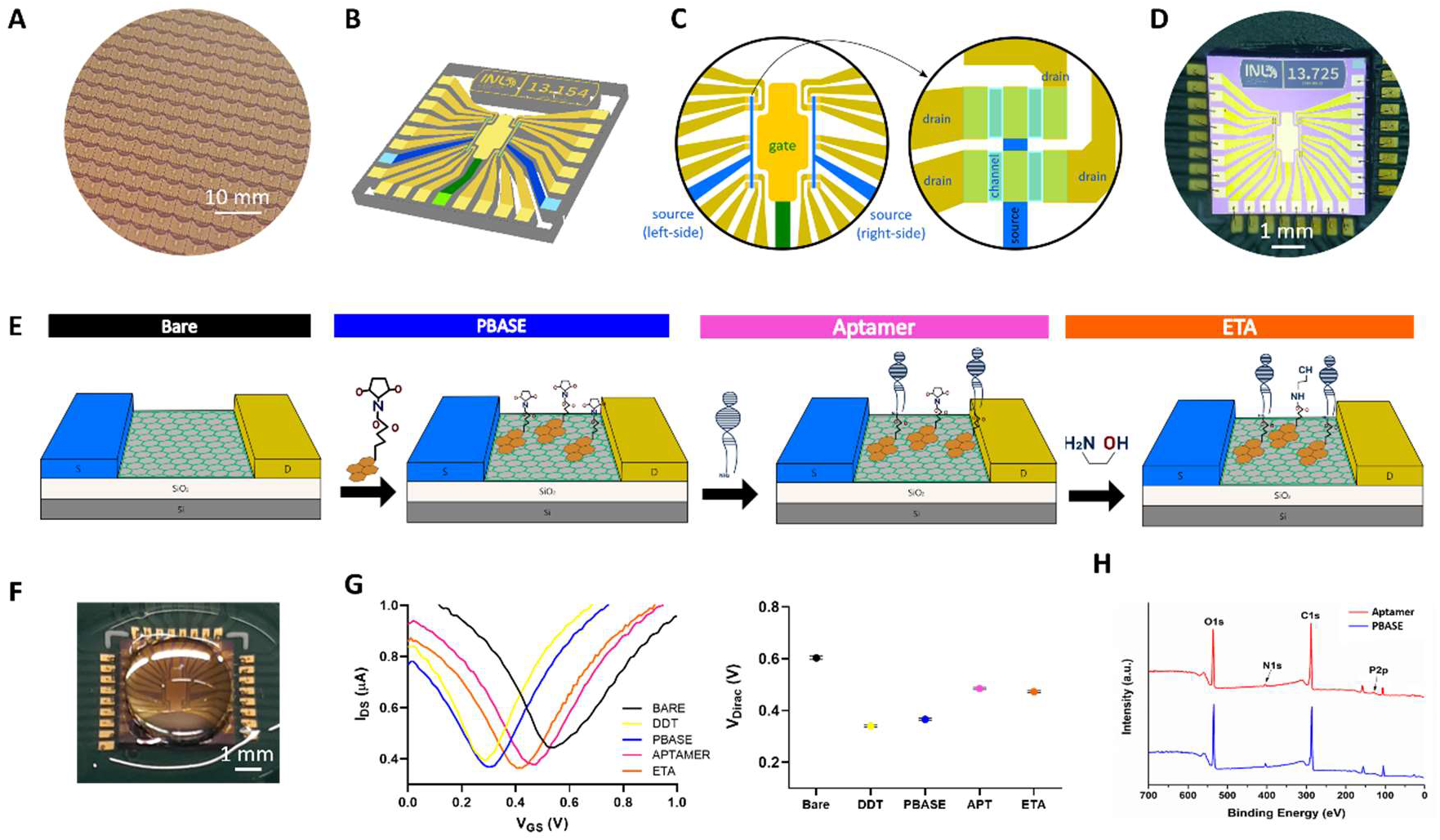
Graphene aptasensor multitransistor arrays (gMTAs) for dopamine detection. (A) Detail of wafer containing 784 fabricated gMTAs. (B) Schematic illustration of one gMTA chip with 20 electrolyte-gated graphene field-effect transistors (EG-gFETs) and respective interconnect lines and pads. (C) Schematic illustration of a gMTA’s sensor area with EG-gFETs sharing a co-planar integrated gate electrode (golden central region and green interconnect line), individual drain electrodes (yellow contacts and interconnect lines), and 2 common source electrodes for every 10 groups of transistors (blue contacts and interconnect lines) (left). Detail of 4 transistors with 4 graphene channels (light blue) connecting 4 independent drain electrodes (yellow) and a common source electrode (dark blue) (right). (D) Photograph of one gMTA wire-bonded to a custom-made PCB for electronics interfacing. (E) Schematic of graphene biofunctionalization process for each EG-gFET in the gMTAs: the exposed bare graphene channel (left) is initially passivated with a pyrene-derived crosslinker (PBASE) (center-left), followed by the addition of a dopamine-specific DNA aptamer that binds to PBASE (center-right) and of ethanolamine (ETA) (right) for blocking unreacted PBASE (N.B. schematic not to scale). (F) Photograph of a gMTA with a phosphate-buffered saline (PBS) droplet on top of the sensor area. (G) Representative transfer curves from one EG-gFET as measured in 1 × PBS after each biofunctionalization step (left). Average value of the charge neutrality point (V_DIRAC_) after each functionalization step for 200 transistors (data is mean ± sem) (right). (H) X-ray Photoelectron Spectroscopy (XPS) survey for graphene samples with PBASE (blue) and PBASE + aptamer (red).

### 2.2. Biofunctionalization of graphene surface for gMTA aptasensor

The EG-gFETs’ graphene channel was functionalized with a dopamine-specific biorecognition probe to guarantee selective dopamine detection, and all remaining exposed surfaces in the device were passivated. A non-covalent biofunctionalization strategy was implemented for immobilizing a short-strand dopamine-specific DNA aptamer (Fig 1E). This approach allows simultaneous sensitive transduction and effective screening within the Debye length. The gate electrode was initially passivated with a thin self-assembled monolayer (SAM) of dodecanethiol (DDT) to prevent adsorption onto the gold surface of any solution molecules. Functionalization of the graphene surface was then achieved with 1-Pyrenebutyric acid N-hydroxysuccinimide (PBASE) that non-covalently binds to graphene through *π*–*π* stacking of aromatic side chains [37] (Fig 1E). The use of non-covalent immobilization preserves graphene’s hyper-conjugated aromatic structure and exceptional electronic mobility [21], which is crucial for developing ultrasensitive biosensors. The succinimidyl ester group of PBASE then remains available to form a covalent bond with an amine-terminated biorecognition probe via nucleophilic substitution [16]. This approach was previously used to successfully immobilize DNA probes in graphene-based biosensors [35], [22]. For selective detection of dopamine, a DNA aptamer previously shown to have a high affinity for dopamine, with a binding constant of 0.25 μM [38], [39], was immobilized in the EG-gFETs by binding to PBASE ester group via a 5’ extremity amine-terminated modifier (Fig 1E). Finally, ethanolamine (ETA) was used for blocking any remaining unreacted PBASE after aptamer binding, further reducing potential sources of non-specific binding and detection (Fig 1E). Combining a DNA aptamer for target biorecognition and the EG-gFETs for transduction forms the base of the graphene aptasensor array for ultrasensitive dopamine detection.

To confirm graphene channel biofunctionalization, the shift in the transistors transfer curves, i.e., the transconductance modulation expressed in source-drain current (I_DS_) changes for a particular gate voltage (V_GS_), was assessed after each functionalization step (Fig. 1F,G). Shifts from the baseline transfer curve measurements indicate charge carriers’ redistribution in the graphene channel due to electrostatic potential changes from surface modification. Measuring the gMTAs EG-gFETs transconductance in PBS produced ambipolar “V”-shaped transfer curves, typically reported for graphene FETs [17], [22], [40] (Fig. 1G). This transfer curve, symmetric about the charge neutrality point or Dirac point (V_DIRAC_), with a left branch (V_GS_ < V_DIRAC_) representing the excursion of the electrochemical potential (μ) in the valence band (p-branch) and a right branch (V_GS_ > V_DIRAC_) representing the excursion of μ in the conduction band (n-branch), is very distinct from semiconductor-based FETs and demonstrates the bipolar character of graphene transistors [12]. I_DS_ is at its minimum at the Dirac point, and tracking the V_DIRAC_ value can be used as a proxy of transfer curve shifts (Fig. 1G). In the as-fabricated EG-gFETs, V_DIRAC_ was observed at positive V_GS_ (Fig. 1G), a consequence of unintentional p-doping during the cleanroom lithographic processes. In all transistors used, the first measurement with the exposed graphene channel in PBS, i.e., before any biofunctionalization step, showed V_DIRAC_ between 0.5 and 0.6 V. Gate electrode passivation with DDT produced a sharp negative shift of V_DIRAC_, of −246 ± 51 mV (Fig. 1C). This shift is attributable to the formation of a SAM covering the gold electrode, creating an excess of positive charges in the solution from the dipole moment reorientation of the alkanethiols [24]. The addition of the crosslinker PBASE produced a slight positive shift of V_DIRAC_ of 23 ± 23 mV (Fig. 1G) as PBASE leads to electron withdrawal (p-doping) [24], [37]. However, the addition of the DNA aptamer induced a sharp positive shift of V_DIRAC_ of 113 ± 31 mV (Fig. 1G). This shift is explained by the DNA’s negatively charged phosphate backbone-induced positive charge in the graphene channel. The final step in the functionalization process, blocking free PBASE with ETA, produces a slight negative shift of V_DIRAC_ of −12 ± 11 mV (Fig. 1G). The final value of V_DIRAC_ for each functionalized EG-gFET in the gMTA, i.e., the gMTA aptasensor, was then used as the baseline for dopamine detection experiments.

Although transfer curve measurements are helpful to track and compare the success of the biofunctionalization process across different gMTAs, the aptamer immobilization process was further assessed with X-ray photoelectron spectroscopy (XPS) by comparing experimental peak parameters for O 1s, C 1s, N 1s, P 2p from graphene samples incubated with PBASE and samples incubated with PBASE followed by the aptamer (Fig. 1H, Table 1). The high-intensity C 1s peak in the PBASE sample comes from the graphene present on the substrate, whereas the O 1s and N 1s peaks come from PBASE’s ester group. PBASE also contributes with carbon but only marginally when compared with graphene. The small atomic percentage of P 2p in the PBASE sample is attributed to the silicon wafer. In the PBASE + aptamer sample, there is a visible increase in O 1s, C 1s, and P 2p atomic percentages relative to the PBASE sample. The O 1s and C 1s increases are attributed to the nucleobases and the sugar units that form nucleotides in DNA. The significant P 2p atomic percentage increase is also a distinctive DNA signature since each nucleotide has a phosphate group that forms a phosphate backbone in DNA. Further analysis of the peaks of interest and the graphene-PBASE-aptamer bond structure can be found in the Supporting Information (Fig. S2).

**Table 1.**
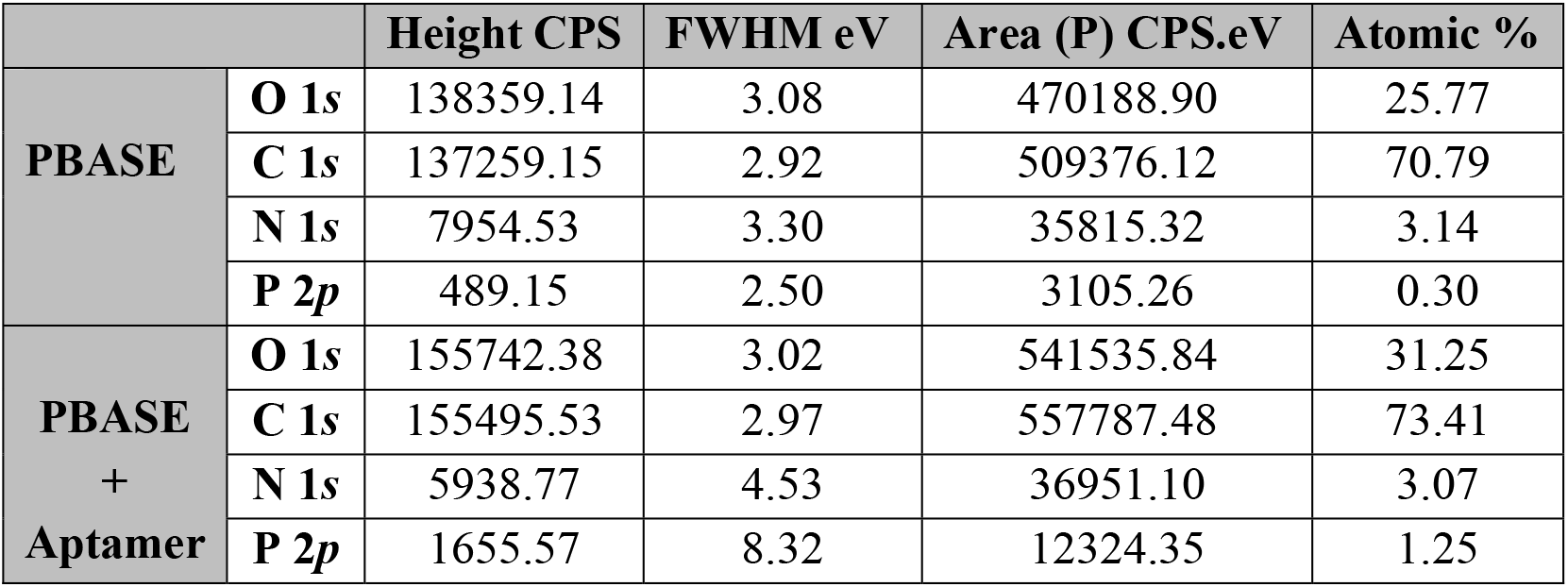
XPS peak parameters for O 1s, C 1s, N 1s, and P 2p for PBASE and PBASE + aptamer on graphene substrates.

|  |  | Height CPS | FWHM eV | Area (P) CPS.eV | Atomic % |
| --- | --- | --- | --- | --- | --- |
| **PBASE** | **O 1s** | 138359.14 | 3.08 | 470188.90 | 25.77 |
|  | **C 1s** | 137259.15 | 2.92 | 509376.12 | 70.79 |
|  | **N 1s** | 7954.53 | 3.30 | 35815.32 | 3.14 |
|  | **P 2p** | 489.15 | 2.50 | 3105.26 | 0.30 |
| **PBASE**<br>**+**<br>**Aptamer** | **O 1s** | 155742.38 | 3.02 | 541535.84 | 31.25 |
|  | **C 1s** | 155495.53 | 2.97 | 557787.48 | 73.41 |
|  | **N 1s** | 5938.77 | 4.53 | 36951.10 | 3.07 |
|  | **P 2p** | 1655.57 | 8.32 | 12324.35 | 1.25 |

### 2.3. Dopamine detection in vitro

For initial validation of the gMTAs ability to detect dopamine in ultra-low concentrations, as well as to establish calibration curves, *in vitro* experiments were performed in undiluted phosphate-buffered saline (1 × PBS) and artificial cerebrospinal fluid (1 × aCSF) (Fig. 2A). Dopamine prepared from stock solution was added to these solutions in increasing concentrations from zM (10^−20^) to nM (10^−9^). The selectivity of the gMTA aptasensor against dopamine synthesis molecules and biological interferents was also assessed.

**Figure 2.**
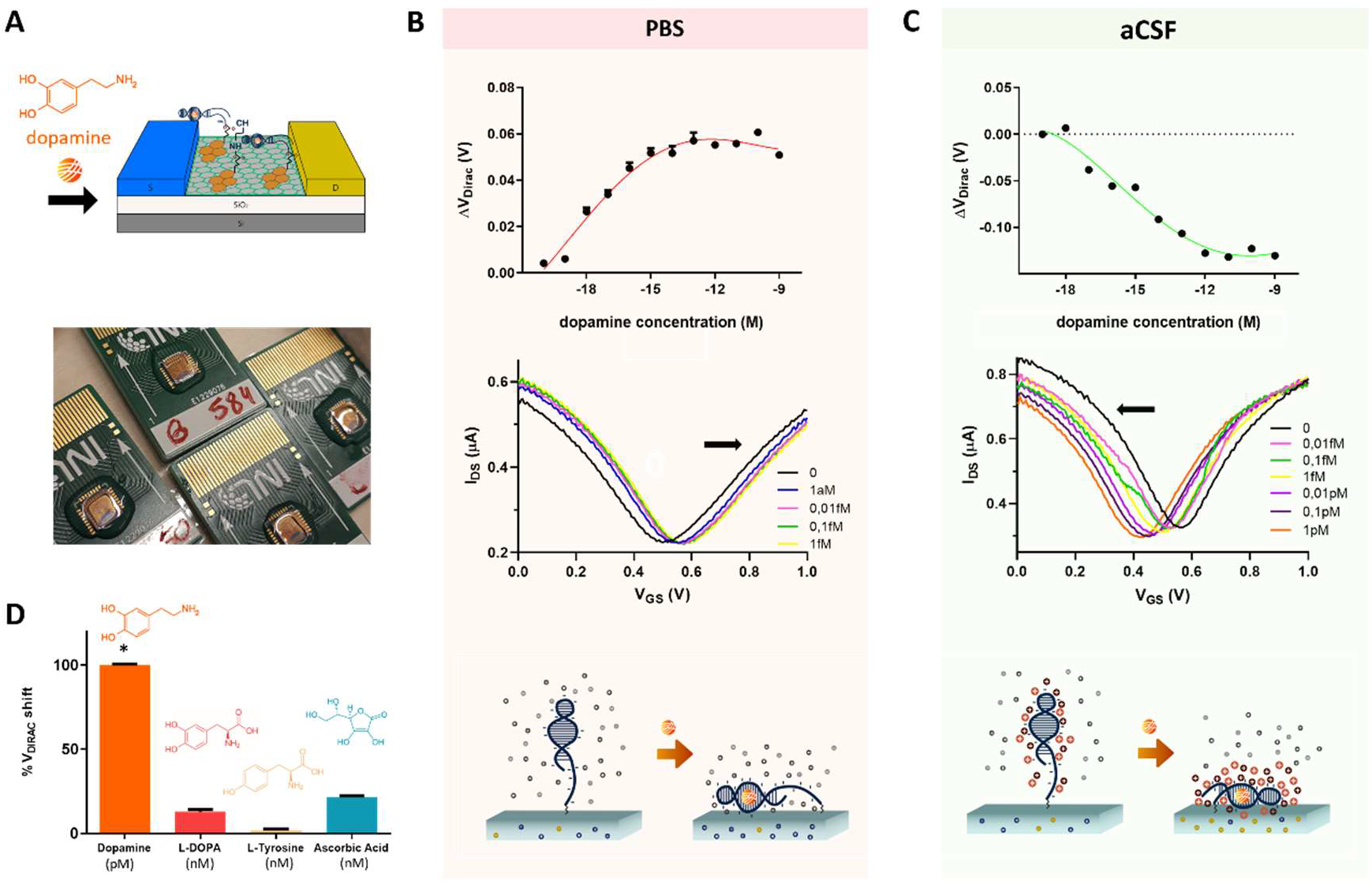
Dopamine detection *in vitro* with gMTAs. (A) Schematic illustration of aptamer structure reorientation close to an EG-gFET graphene channel upon dopamine binding (top). Photograph of four gMTAs incubating 20μL samples containing dopamine in undiluted phosphate-buffered saline (1 × PBS) (bottom). (B) Calibration curve for dopamine detection in 1 × PBS (data is mean ± sem, with 4^th^ order polynomial line fit) (top). Representative transfer curve shifts as a function of increasing dopamine concentrations for the linear detection range of one EG-gFET in 1 × PBS. V_DIRAC_ moves towards positive gate voltages (V_GS_) (black arrow indicates the direction of V_DIRAC_ shifts) (middle). Illustration of the hypothesized reorientation of the aptamer’s backbone negative charges close to the gFETs upon dopamine binding in 1 × PBS, leading to electrostatic repulsion in the graphene channel. (C) Calibration curve for dopamine detection in undiluted artificial cerebral spinal fluid (1 × aCSF) (data is mean ± sem, with 4^th^ order polynomial line fit) (top). Representative transfer curve shifts as a function of increasing dopamine concentrations for the linear detection range of one EG-gFET in 1 × aCSF, with V_DIRAC_ moving towards negative gate voltages (V_GS_) (black arrow indicates the direction of V_DIRAC_ shifts) (middle). Illustration of the hypothesized reorientation of the aptamer’s structure in aCSF upon dopamine binding with the increased attraction of positive charges closer to the graphene channel, leading to negative shifts of V_DIRAC_ (bottom). (D) Comparative responses of gMTAs to 1 pM dopamine, 1 nM L-DOPA, 1nM L-tyrosine and 1 nM ascorbic acid in 1 × PBS (normalized data is mean + sem; One-way ANOVA, p<0.0001).

#### 2.3.1 Dopamine detection in PBS

A calibration curve for the gMTAs response (ΔV_DIRAC_) to dopamine was obtained by serially incubating solutions of increasing dopamine concentrations in undiluted phosphate-buffered saline (1 × PBS) (Fig. 2B). V_DIRAC_ measurements were offset-corrected by 19.9 mV, which was the mean ΔV_DIRAC_ obtained for blank samples, i.e., samples prepared from dilution of dopamine stock solution but not containing dopamine molecules (Supporting Information, Fig. S3) Adding dopamine to the gMTAs led to a significant V_DIRAC_ shift of approximately 26 ± 1 mV for a concentration as low as 1 aM (10^−18^) (Fig. 2B), a record limit-of-detection (LOD) for dopamine. The intrinsic variance of the gMTAs’ measurements was also calculated by assessing their response to blank samples over time (Supporting Information, Fig. S4). The obtained coefficient of variation (CV) for ΔV_DIRAC_ was 1.13%, which is indicative of the gMTA’s high stability in the presence of solution ions, and well below the observed 26 ± 1 mV V_DIRAC_ shift obtained for 1 aM (10^−18^) dopamine concentration. From 1 aM (10^−18^) dopamine concentration, gMTAs presented a linear detection range up to 1 fM (10^−15^), with linear increases of V_DIRAC_ as a function of dopamine concentration with a 9.5 mV/decade sensitivity (Fig. 2B). The observed positive shifts in V_DIRAC_ with increasing concentrations of dopamine are hypothesized to occur due to reorientation of the negative charges of the DNA aptamer phosphate backbone near the EG-gFETs’ channels upon dopamine binding [41] (Fig. 2B). These charges, acting as counter-ions, would raise the energy of the graphene electron bands, effectively shifting the electrochemical potential, μ, towards the valence band. Consequently, the charge neutrality point (V_Dirac_) is found at more positive gate voltages, required to bring μ back to the Dirac point.

The dopamine LOD of 1 aM (10^−18^) obtained with our gMTAs is 3 orders of magnitude lower than the lowest LOD ever reported for any dopamine sensor, currently at 0.5 fM (10^−15^) [8], [9], and several orders of magnitude lower than the LOD attained with the majority of previous methodologies, as summarized in the Supporting Information (Table S1). Additionally, the sensors currently presenting the lowest dopamine LOD, although based on organic field-effect transistors [12] or graphene electrodes [13], rely on voltammetric electrochemical measurements or require the addition of labels/reporters. The gMTA aptasensor is a label-free detection device, requiring low operational voltage with high transconductance due to the high gate capacitance from the electrical double layers (EDLs) at the graphene-electrolyte and electrolyte-gate interfaces. The record-breaking sensitivity of our sensor results from a combination of factors: (i) the EG-gFETs high 2D conductance, deriving from CVD graphene’s single-layer high electronic mobility and relatively high carrier density [14], [16]; (ii) the cleanroom fabrication process carefully developed to preserve graphene’s electronic properties, while simultaneously passivating all other device areas [36]; (iii) the graphene transistor channel direct exposure to the liquid medium containing the target and the EDLs formation [17], [42]; (iv) the aptamer’s affinity to dopamine and its ability to operate within the Debye length [38], [39], [43].

#### 2.3.2 Dopamine detection in aCSF

To further validate the ultrasensitivity of our gMTAs in a more complex *in vitro* medium with physiologically high ionic strength, calibration curves for dopamine detection in undiluted artificial cerebrospinal fluid (1 × aCSF) were acquired (Fig. 2C). Although aCSF does not contain other molecules found in biological CSF, such as amino acids, proteins, and hormones, it closely matches biological CSF’s electrolytic profile and osmolarity. The observed LOD in 1 × aCSF was 10 aM (10^−17^) (Fig. 2C), which is one order of magnitude higher than that observed in 1 × PBS. This difference was expected because the Debye length is lower in aCSF when compared with PBS due to a higher ionic strength, at the same pH, attributable to additional divalent cations in the solution. Nevertheless, and although most dopamine sensors are not tested in aCSF, the observed LOD is still 3 orders of magnitude lower than the previously reported LOD for a dopamine sensor in aCSF at 10 fM (10^−14^) [39]. The dynamic detection range is also broader in aCSF compared with PBS, going up to 100 nM (10^−11^) with a 22 mV/decade sensitivity (Fig. 2C).

Contrary to the calibration curve in 1 × PBS, adding dopamine to the gMTAs in 1 × aCSF led to negative shifts of V_DIRAC_, which means that the graphene electrochemical potential moved up in energy relative to the density of states (n-doping). The sign of V_DIRAC_ shifts depends on several mechanisms involving interactions between the probe and the target and between the electrolytic solution and the probe and target [16]. Thus, one possible interpretation for the observed negative shift is that the secondary structure of dopamine aptamer is different in aCSF, when compared with that in PBS, which, consequently, brings positively charged molecular regions towards the graphene channel upon dopamine binding (Fig. 2C). These results lend further support to previous observations reporting potential alterations of the secondary structure of this aptamer in the presence of Ca^2+^ and Mg^2+^, which are present in both CSF and aCSF [38], [41]. The excess of positive charges near the graphene channel upon dopamine binding may also explain the significant increase of the dynamic detection range in aCSF, because each detection event may add multiple positive charges within the Debye length.

#### 2.3.3 Selectivity assessment

Evaluating the selectivity of novel biosensors is paramount for successful real-world applications with biological samples where the target molecule is mixed with various other molecules. Brain dopamine or dopamine in brain-derived samples, such as biological CSF, occurs at minute concentrations and is mixed with other neurotransmitters, amino acids, and proteins. Thus, to test the selectivity of the gMTAs, their response to dopamine was compared with the response to molecules in the dopamine synthesis pathway, including L-Dopa and L-Tyrosine, which are chemically similar to dopamine, and ascorbic acid, a ubiquitous biological interferent typically occurring in higher concentrations than neurotransmitters. The latter is also involved in norepinephrine synthesis from dopamine, and therefore likely to co-occur with dopamine in the brain [27]. The response of the gMTAs to the other tested molecules, even when present in high concentrations (1 nM), was negligible compared with the response to low concentrations of dopamine (1 pM) (Fig. 2D), confirming the aptamer’s specificity and high affinity to dopamine. The response to other monoamine neurotransmitters, such as serotonin and norepinephrine, was not tested, but a previous report has demonstrated that this aptamer has reduced affinity for those neurotransmitters [39].

### 2.4. Dopamine detection in an animal model of Parkinson’s Disease

Biosensors developed and optimized based on *in vitro* assays tend to underperform in biological samples, losing sensitivity and selectivity. This underperformance, also observed in previous ion-sensitive dopamine biosensors [8], [9], is usually due to a high number of interferent molecules and ions typically not found in optimized buffers and the reduction of the Debye length due to the increased shielding effect of the biological solution’s additional counter-ions [44]. Thus, to evaluate the performance of our gMTAs in a physiological scenario, dopamine detection experiments were performed in biological CSF and brain homogenate samples.

A reserpine-induced mouse model of Parkinson’s Disease (PD) was used for these experiments. This animal model has been instrumental in elucidating the role of abnormal dopamine levels in PD symptomatology and used to develop critical therapeutics for PD, such as L-DOPA administration [45], [46]. Reserpine is an irreversible and non-selective inhibitor of the vesicular monoamine transporters VMAT1 and VMAT2, and its systemic administration impairs monoamine uptake and storage in neuronal cells leading to the rapid depletion of brain dopamine from neuronal synapses [47]–[49]. This depletion leads to impaired motor function resulting in behavioral symptoms similar to PD, caused by the loss of dopaminergic neurons in the human brain [48], [49]. As expected, 8h postadministration, all our reserpine-treated mice displayed severe akinesia, postural instability, and tremors, while controls (vehicle-treated mice) displayed standard motor and exploratory behavior.

#### 2.4.1 Dopamine detection in biological CSF

To test if our gMTAs could detect relevant small changes in dopamine concentration from small working volume biological samples, their ability to differentiate CSF samples obtained from control and parkinsonian (resperine-treated) mice was assessed (Fig. 3A). CSF was extracted by *cisterna magna* puncture [50], a method similar to subarachnoid puncture used in humans for clinical diagnostic [51]. Pairs of 2μL CSF samples obtained from reserpine-treated (dopamine-depleted) and control animals were incubated sequentially, in this order, on each gMTA. A five-fold response difference was observed when comparing V_DIRAC_ shifts of samples from dopamine-depleted animals (reserpine, mean ΔV_DIRAC_: 9 ± 8 mV) with those of control animals (mean ΔV_DIRAC_: 51 ± 11 mV) (Fig. 3A). Of note, CSF samples were directly incubated in the gMTAs after extraction without the need for any sample preparation, and measurements were acquired after 10 minutes of incubation only. This result is of great importance for developing fast-acting point-of-care devices. Additionally, the gMTAs displayed significantly higher sensitivity than previously developed biosensors for dopamine detection in CSF while requiring significantly lower sample working volumes [52]–[54]. The ability of our gMTAs in detecting minimal changes in dopamine concentration in biological samples is remarkable, as detecting clinically relevant dopamine alterations in PD patients, even before motor symptoms arise, is a highly sought-after goal for the development of earlier and improved diagnostics for PD.

**Figure 3.**
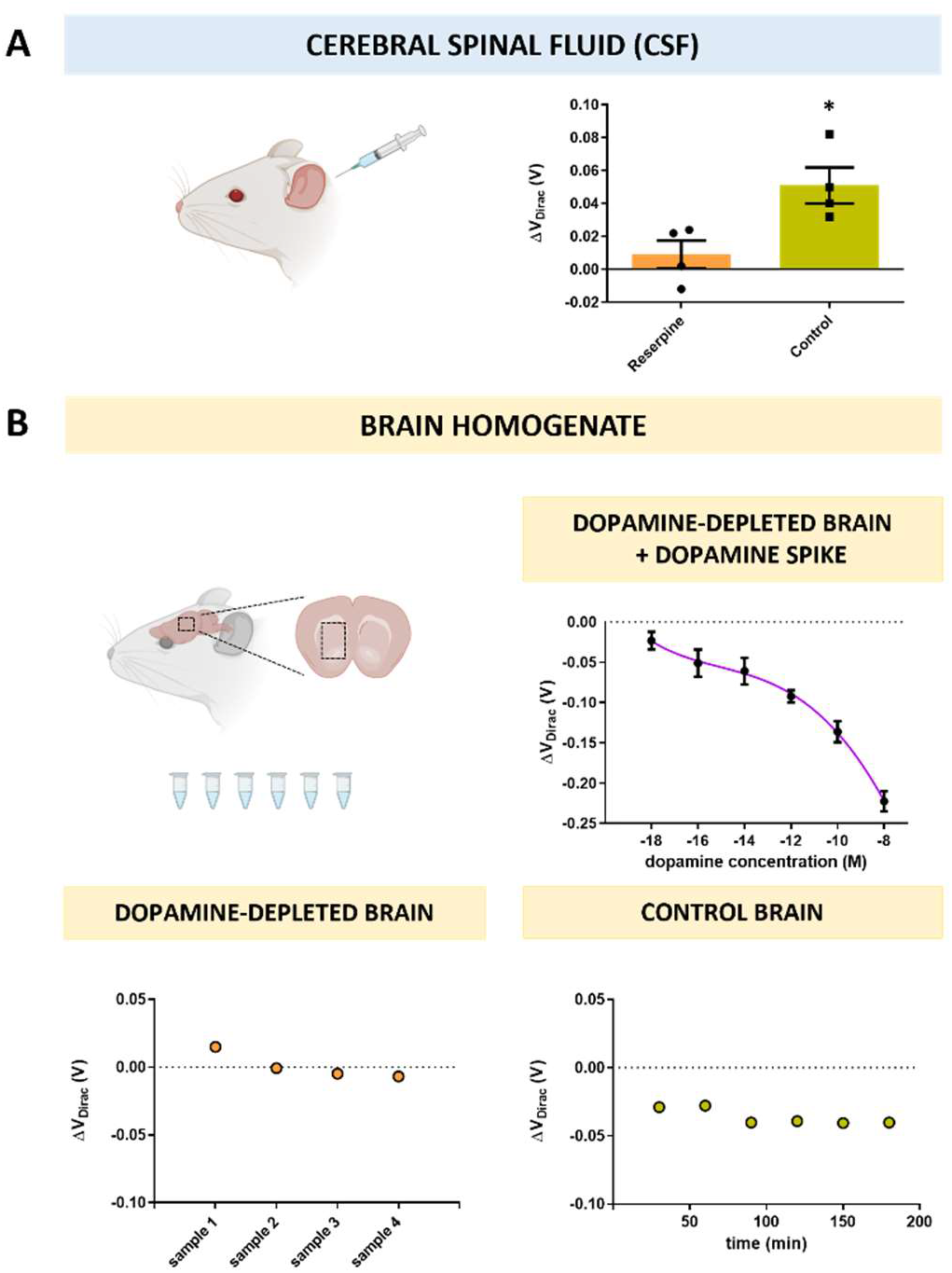
Dopamine detection with gMTAs in biological samples from a mouse model of Parkinson’s Disease (PD). (A) Response of gMTAs to biological cerebral spinal fluid (CSF) samples obtained from PD (reserpine) and control mice (data is mean ± sem; paired t-test, p<0.05). (B) gMTAs response to dopamine-depleted brain homogenate samples obtained from PD mice spiked with increasing dopamine concentrations (top) (data is mean ± sem, with 4^th^ order polynomial line fit). gMTAs response to sequentially incubated dopamine-depleted brain homogenate samples obtained from PD mice (bottom-left). gMTAs response to brain homogenate sample obtained from control mice incubated continuously for 3 hours.

#### 2.4.2 Dopamine detection in brain homogenate

Finally, to test our gMTAs’ dopamine detection in a media that more closely mimics the complexity of the brain’s extracellular space, homogenate brain samples from reserpine-treated (dopamine-depleted) animals were collected and spiked with increasing dopamine concentrations, prepared from stock solution, from aM (10^−18^) to nM (10^−8^) (Fig. 3B). The gMTAs still displayed high sensitivity for dopamine detection, overcoming sensitivity losses typically observed in such complex media (Fig. 3B). The observed LOD was 1 aM (10^−18^), and a wide dynamic range from 1 aM (10^−18^) to 100 μM (10^−8^) was obtained (Fig. 3B).

Two additional control experiments were performed to confirm the selective detection of dopamine in brain homogenates by the gMTAs. First, samples from reserpine-treated animals without dopamine spiking were incubated in the gMTAs to confirm that the previously observed changes in transconductance were due to the selective detection of dopamine and not other molecules present in the brain homogenates (Fig. 3B). As expected, no significant V_DIRAC_ shifts from baseline values were observed between samples, denoting the absence of dopamine detection (Fig. 3B). Then, one homogenate sample pooled from 2 control animals’ brains was continuously incubated in one gMTA for 180 minutes, and measurements were taken every 30 minutes. This experiment was performed to confirm that the observed significant shifts in V_DIRAC_ from the dopamine spiked samples were due to the presence of dopamine and not a time-dependent variation in transconductance. A sharp 29 mV V_DIRAC_ shift was observed for the first measurement, after 30 minutes incubation, attributed to the presence of dopamine in the control brains (Fig. 3B). Then, no significant V_DIRAC_ shifts between measurements were observed for the remaining 150 minutes (Fig. 3B). The fact that dopamine can be reliably detected in ultra-low concentrations in samples resembling the brain’s extracellular space with the gMTAs paves the way for novel *in vivo* biosensors requiring localized dopamine measurements in a complex biological scenario.

## 3. Conclusions

The graphene aptasensor multitransistor array (gMTA) for dopamine detection proposed here, combining an array of graphene field-effect transistors with a selective DNA aptamer, achieved ultrasensitive and stable dopamine detection in various media from physiological ionic-strength buffers to complex biological samples. Notably, the gMTAs detected dopamine changes in small working volume CSF samples from a Parkinson’s Disease (PD) animal model, which can pave the way for novel point-of-care diagnostic devices to detect abnormal levels of dopamine in PD, and in other dopamine-dependent brain disorders such as Alzheimer’s Disease, and substance addiction disorders. The use of numerous EG-gFET aptasensors in an array configuration further allows future multiplexed measurements of different biomarkers in one single gMTA through the localized biofunctionalization of individual EG-gFETs. Lastly, the fabrication at the wafer level lends itself to developing different array configurations, including with higher EG-gFETs count, which may be relevant for a high-throughput assessment of localized dopamine release. Increased sensors’ spatial density and tracking independent localized measurements with single-cell resolution from each EG-gFET or subsets of transistors in the arrays are future features that can be realized without significant changes to the overall fabrication process. This possibility is especially relevant for fundamental neuroscience or pharmaceutical studies of brain disorders requiring measurements from *ex vivo* brain slices from animal models or human patients’ IPSC-derived cell cultures and *in vivo* monitoring in intact brains. As the proposed sensor is further tested in other biologically relevant samples and in *in vivo* scenarios, it can ultimately help elucidate our understanding of the brain and promote the development of improved diagnostics and therapeutics for brain disorders.

## 4. Materials and Methods

### 4.1 Reagents and Materials

3,4-dihydroxyphenethylamine hydrochloride (Dopamine hydrochloride), 3,4-Dihydroxyl-L-phenylalaline (L-DOPA) (98% TLC), L-Tyrosine (98% HPLC), L-Ascorbic acid (99%), Phosphate-buffered saline (PBS) tablets, N,N-Dimethylformamide (DMF) (99.9% HPLC), 1-Pyrenebutynic acid N-hydroxy-succinimide ester (PBASE) (95%), 1-Dodecanethiol (DDT) (98%), Ethanolamine (ETA) (98%), Sodium Chloride (NaCl), Potassium Chloride (KCl), Monosodium Phosphate (NaH2PO4), Sodium Bicarbonate (NaHCO3), Glucose, Calcium Chloride Dihydrate (CaCl2.2H2O), Acetic Acid Glacial, and Poly(methyl(meth)acrylate) (PMMA) (15k M.W.) were purchased from Sigma-Aldrich. Acetone (99.5%), Ethanol (99.8%), and 2-Propanol (99,8% GC) were purchased from Honeywell. Magnesium Sulfate Heptahydrate (MgSO4.7H2O) was purchased from Merck. Photoresist AZ1505 (AZ) was purchased from MicroChemicals GmbH. RTV silicone elastomer (3140, Dowsil) and superglue (Loctite) were acquired from Farnel. DNA aptamer (5′-CGACGCCAGTTTGAAGGTTCGTTCGCAGGTGTGGAGTGACGTCG-3′) with a 5′ C6-amino link modification was synthesized by STABvida. Reserpine was acquired from Biogen. MilliQ water used in all experiments had a resistivity higher than 18 MΩ cm at 25°C.

### 4.2 Graphene multitransistor array fabrication

Graphene multitransistor arrays (gMTAs) were fabricated by our previously published optimized process for high-yield wafer-scale fabrication of electrolyte-gated graphene field-effect transistors (EG-gFETs) with an integrated gate electrode and improved multilayer dielectric passivation for biosensing applications [36]. Summarized methods are described below.

#### 4.2.1 Single-layer graphene growth

Briefly, single-layer graphene (SLG) was grown by thermal chemical vapor deposition (CVD) on 25 μm thick high-purity (99.999% purity) copper (Cu) foils in a three-zone quartz tube furnace (EasyTube ET3000, CVD Corp.). The 10 × 10 mm^2^ Cu substrates were initially treated with a mixture of FeCl_3_, HCl, and deionized water (DI) for 1 minute in ultrasound and then placed in the furnace. The system was evacuated to approximately 2 mTorr and then filled with 250-sccm Argon (Ar, 99.999% purity) and 60-sccm Hydrogen (H_2_, 99.999% purity) gas mixture. Once the growth temperature and pressure were reached, methane (0.5 sccm), the carbon precursor, was introduced into the chamber. The growth was carried out at 1040 °C at 6 Torr for 25 minutes. After growth, SLG was protected by poly(methyl(meth)acrylate) (PMMA), and plasma ashing was performed to remove the graphene from the backside of the substrate. Raman spectroscopy was used to assess the quality of grown graphene (Supporting Information, Fig. S1)

#### 4.2.2 Wafer-level fabrication of electrolyte-gated graphene field-effect transistors

Briefly, a 200 mm silicon (Si) wafer with 100 nm of thermal oxide was used as a substrate. The wafer was sputter-coated with Chromium (Cr) (3 nm) as an adhesion layer, Gold (Au) (35 nm) as the conductive layer, and an alumina (Al_2_O_3_) (20 nm) capping. The source, drain, and gate electrodes were patterned by optical lithography and etched by ion milling. A sacrificial layer (TiW, 5 nm; AlSiCu, 100 nm; TiWN, 15 nm) was sputtered and patterned via lift-off, exposing the channel region and the source and drain electrodes for graphene transfer. Previously grown SLG, as described above, was transferred onto the wafer, patterned with optical lithography, and dry-etched with oxygen plasma. The sacrificial layer was then removed by wet etch. A protective layer (Ni, 10 nm; AlSiCu 30 nm; TiWN 10 nm) was sputtered and patterned by sonication-free lift-off to work as a stopping layer for reactive ion etching (RIE) at the graphene channel and gate electrode. A 250 nm multi-stack passivation layer of SiO_2_ and Si_3_N_4_ was grown by plasma-enhanced CVD and patterned by RIE. Finally, the stopping layer was dry-etched to expose the graphene channel and gate electrode.

The EG-gFETs were characterized electrically at the wafer level with an automated probe station by measuring the resistance between each transistor’s source and drain electrodes at a fixed voltage of 1 mV or a fixed current of 1 μA. gMTAs with transistors with resistance above 2.5 MΩ were discarded from further experiments.

The wafer was finally coated with photoresist for protection and diced into individual 4.5 × 4.5 mm gMTA chips. The chips were then glued onto custom-designed printed circuit boards (PCB) for electronic interfacing, and their interconnection pads were wire-bonded with gold wires to the PCB pads. Finally, the wires and connection pads were protected with a silicone elastomer.

### 4.3 Graphene transistors functionalization with dopamine-specific aptamer

EG-gFETs in each gMTA chip were first incubated for 4 hours in 2 mM 1-Dodecanethiol (DDT, 2mM in ethanol) for gate passivation, then cleaned with DI water and dried under N_2_ flow. Then, 20 μL of the pyrene-derived crosslinker 1-Pyrenebutyric acid N-hydroxysuccinimide (PBASE, 10 mM in dimethylformamide, DMF), were added to the gMTAs and incubated for 2 hours in a humid chamber. A 44nt-long DNA aptamer (5’-CGACGCCAGTTTGAAGGTTCGTTCGCAGGTGTGGAGTGACGTCG-3′), previously selected for high affinity to dopamine [39], with a 5′ amino-link termination, was initially diluted in MilliQ water to 20 μM, heated to 95°C for 5 minutes, and then allowed to cool to room temperature. Then, 20 μL of aptamer solution was added to the gMTAs and incubated for 16 hours in a humid chamber in the dark. 20 μL of ethanolamine (ETA, 100 mM in DI water) were incubated in the gMTAs for 30 minutes to bind to PBASE via the amine termination and block any remaining PBASE that did not bind to the aptamers. Finally, gMTAs were rinsed with DI water and dried under N_2_ flow. The transconductance of each EG-gFET aptasensor in the arrays was measured at each step of the functionalization process by applying a source-drain voltage (V_DS_) of 1 mV and measuring source-drain current (I_DS_) in gate-source voltage (V_GS_) sweeps between 0 and 1 V.

### 4.4 X-ray photoelectron spectroscopy (XPS)

X-ray photoelectron spectroscopy (XPS) analysis was performed with an ESCALAB 250Xi system (Thermo Scientific). A thermal p-doped silicon oxide wafer was coated with a photoresist and cut in 1 mm^2^ substrates. Previously grown SLG was transferred to the substrate, after which samples were either incubated with the PBASE crosslinker (for 2 hours) or with PBASE followed by incubation with the dopamine-specific aptamer (for 16 hours). Survey scans were performed from 0 to 1350 eV with a pass energy of 50 eV. High-resolution spectra were completed for carbon, nitrogen, oxygen, and phosphorous. Advantage software (Thermofisher) was used for peak fitting and calculating atomic percentages.

### 4.5 In vitro sample preparation and measurement protocols

Dopamine hydrochloride (3,4-dihydroxyphenethylamine hydrochloride) in powder form was first prepared into a stock solution of 10 mM in either phosphate-buffered saline (PBS) (mM: 137 NaCl, 2.7 KCl, 10 KH_2_PO_4_, 1.8 NaH_2_PO_4_) or artificial cerebral spinal fluid (aCSF) (mM: 119 NaCl, 2.5 KCl, 1.2 NaH_2_PO_4_, 24 NaHCO_3_, 12.5 glucose, 2 MgSO_4_.7H_2_O and 2 CaCl_2_.2H_2_O, 300-310 mOsm/L) at pH 7.2-7.4. Solutions of different dopamine concentrations in 1 × PBS or 1 × aCSF were prepared by diluting the stock solutions from zM (10^−20^) to nM (10^−9^). Baseline transconductance for EG-gFETs in each gMTA aptasensor was measured in PBS or aCSF not containing dopamine by applying a V_DS_ of 1 mV, and sweeping V_GS_ between 0 and 1 V. Following baseline measurements, 20 μL samples of each dopamine concentration in either buffer were incubated in the gMTAs. Between samples, gMTAs were rinsed with DI water, and transconductance measurements were taken in 1 × PBS or 1 × aCSF accordingly, with the same voltage parameters used for baseline acquisition. More than 500 EG-gFETs from over 30 gMTAs provided measurements for the PBS and aCSF calibration curves, with each concentration for each electrolytic buffer being incubated in at least 4 gMTAs (minimum 80 EG-gFETs per concentration per buffer).

L-DOPA (L-3,4-dihydroxyphenylalanine), L-Tyrosine, and ascorbic acid were diluted in 1 × PBS to 1 nM (10^−9^) concentration. Then, 20 μL samples of each solution were incubated in the gMTAs. After DI water rinse, transconductance measurements were taken in 1 × PBS with the same voltage parameters described above. Samples from each solution were incubated in at least 2 gMTAs (minimum 40 EG-gFETs per solution).

### 4.6 Biological samples preparation and measurement protocols

All animal procedures described in this work complied with the European Union Directive 2016/63/EU and the Portuguese regulations and laws protecting animals used for scientific purposes (DL No 113/2013). The Ethics Subcommittee for the Life Sciences and Health of Minho University and the Portuguese National Authority for Animal Health approved this study.

Mice (n=8) were injected with reserpine (5 mg/kg, i.p.) dissolved in 1% glacial acetic acid and diluted in 0.9% saline. Control mice (n=6) were injected with vehicle solution not containing reserpine. 8h post-administration, all mice were anesthetized with avertin (tribromoethanol, 20 mg/mL; 0.5 mg/g, i.p.), placed in a stereotaxic frame for head fixation, and CSF was collected by *cisterna magna* puncture. CSF samples were inspected for blood contamination and immediately frozen in liquid nitrogen and stored at −80°C. Mice were then perfused transcardially with 0.9% saline to remove circulating blood, and the brain was quickly removed. The striatum brain region, on both hemispheres, was dissected, frozen in liquid nitrogen, and stored at −80°C.

For dopamine detection in CSF, frozen samples were thawed on ice, and 2 μL samples were placed directly on the gMTAs. For every gMTA, transconductance measurements were first acquired for a CSF sample from a reserpine-treated animal incubated for 10 minutes. Following rinsing with PBS, a sample from a control animal was incubated in the same gMTA for 10 minutes. The EG-gFETs transconductance was measured by applying a V_DS_ of 1 mV and sweeping V_GS_ between 0 and 1 V.

Previously dissected striatum brain samples were first thawed on ice for dopamine detection in brain homogenate. Samples from two brains (total of 4 hemispheres) were pooled in a single Eppendorf tube with 1 × aCSF (pH 7.4) added at 5 μL/mg for homogenization with a brain tissue homogenizer. Samples were then centrifuged at 14000 rpm, for 15 mins at 4°C. Dopamine hydrochloride diluted in 1 × aCSF was added to 10 μL of homogenate supernatant samples from reserpine-treated animals on a 1:1 volume ratio for final dopamine concentrations ranging from 1 aM (10^−18^) to 10 nM (10^−8^). Each 20 μL sample was incubated for 20 minutes in one gMTA and then rinsed with 1 × PBS. Transconductance measurements were taken before rinsing with the same voltage parameters described above.

As negative controls, 20 μL brain homogenate samples pooled from two reserpine-treated animals diluted in aCSF on a 1:1 volume ratio without dopamine added were serially incubated in one gMTA for 20 minutes. Transconductance measurements and rinsing between samples were performed as described above. One 20 μL brain homogenate sample pooled from two control animals diluted in aCSF on a 1:1 volume ratio was also incubated continuously for 3 hours in one gMTA. Transconductance measurements were taken every 30 minutes without rinsing with the same voltage excursion as above.

### 4.7 Statistical Analysis

Statistical analysis was performed with either GraphPad Prism 9 for Windows (Graphpad Software) or Matlab ver. 2021a (Mathworks). Statistical details can be found in figure legends.

## CRediT authorship contribution statement

**Mafalda Abrantes:** Investigation, Formal Analysis, Data Curation, Visualization, Writing – Original Draft. **Diana Rodrigues**: Investigation. **Telma Domingues:** Methodology. **Siva S. Nemala:** Investigation, Visualization. **Patricia Monteiro**: Validation, Resources, Supervision, Funding Acquisition. **Jérôme Borme:** Methodology, Visualization, Validation, Supervision. **Pedro Alpuim:** Conceptualization, Validation, Resources, Writing – Reviewing and Editing, Supervision, Funding Acquisition. **Luis Jacinto**: Conceptualization, Formal Analysis, Validation, Visualization, Resources, Writing – Original Draft, Writing – Review and Editing, Supervision, Funding Acquisition.

## Acknowledgments

The authors would like to thank Prof. Fátima Cerqueira of the University of Minho and the International Iberian Nanotechnology Laboratory for acquiring the Raman spectra of graphene samples. Rodent figures were created with BioRender.com. This work was funded by “la Caixa” Banking Foundation under the grant agreement LCF/PR/HR21-00410; national funds, through the Foundation for Science and Technology (FCT) - projects UIDB/50026/2020 and UIDP/50026/2020; by FCT project PTDC/MED-NEU/28073/2017 (POCI-01-307 0145-FEDER-028073); and by The Branco Weiss fellowship - Society in Science (ETH Zurich).

## SUPPORTING INFORMATION

**Table S1.**
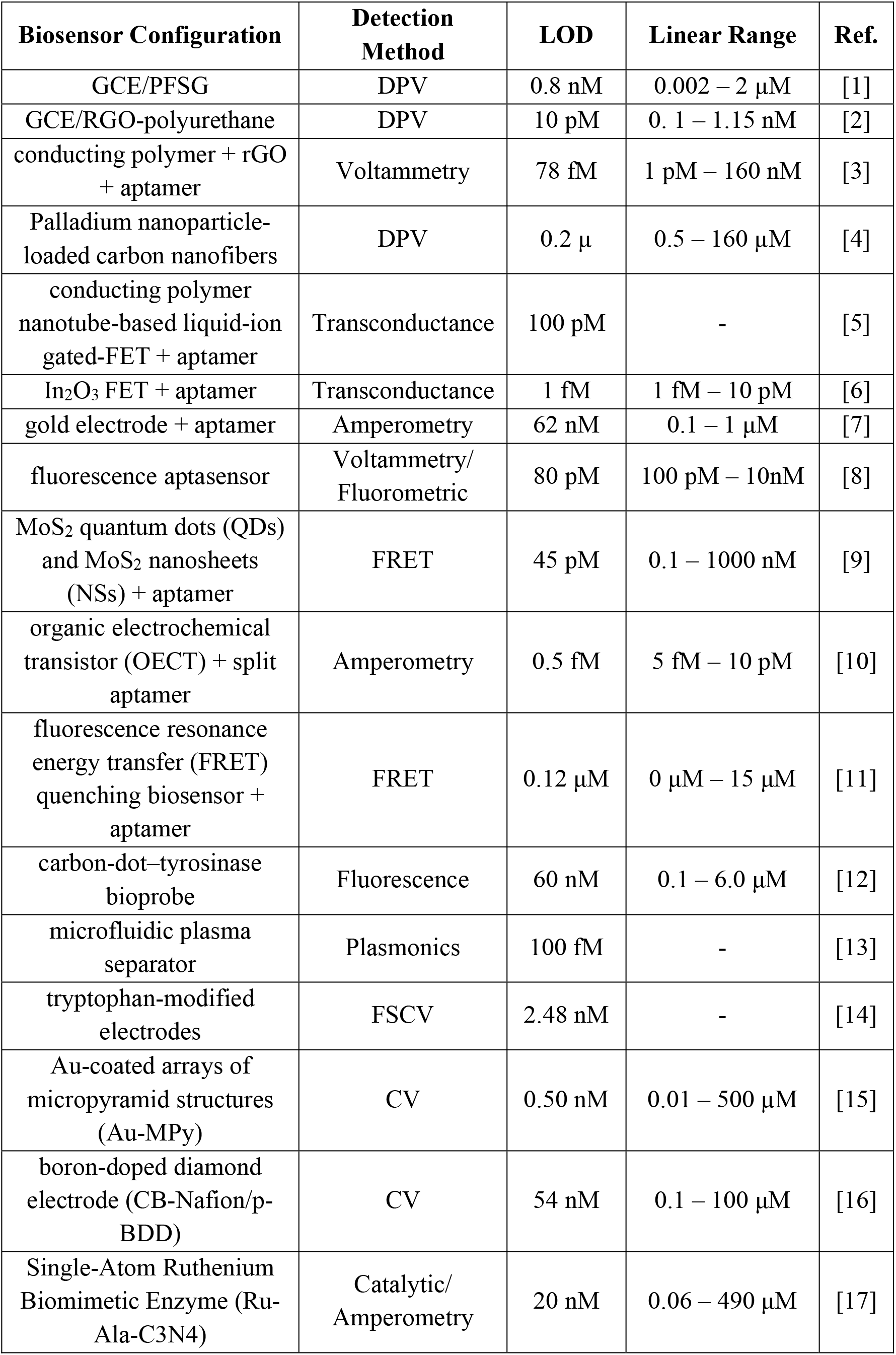
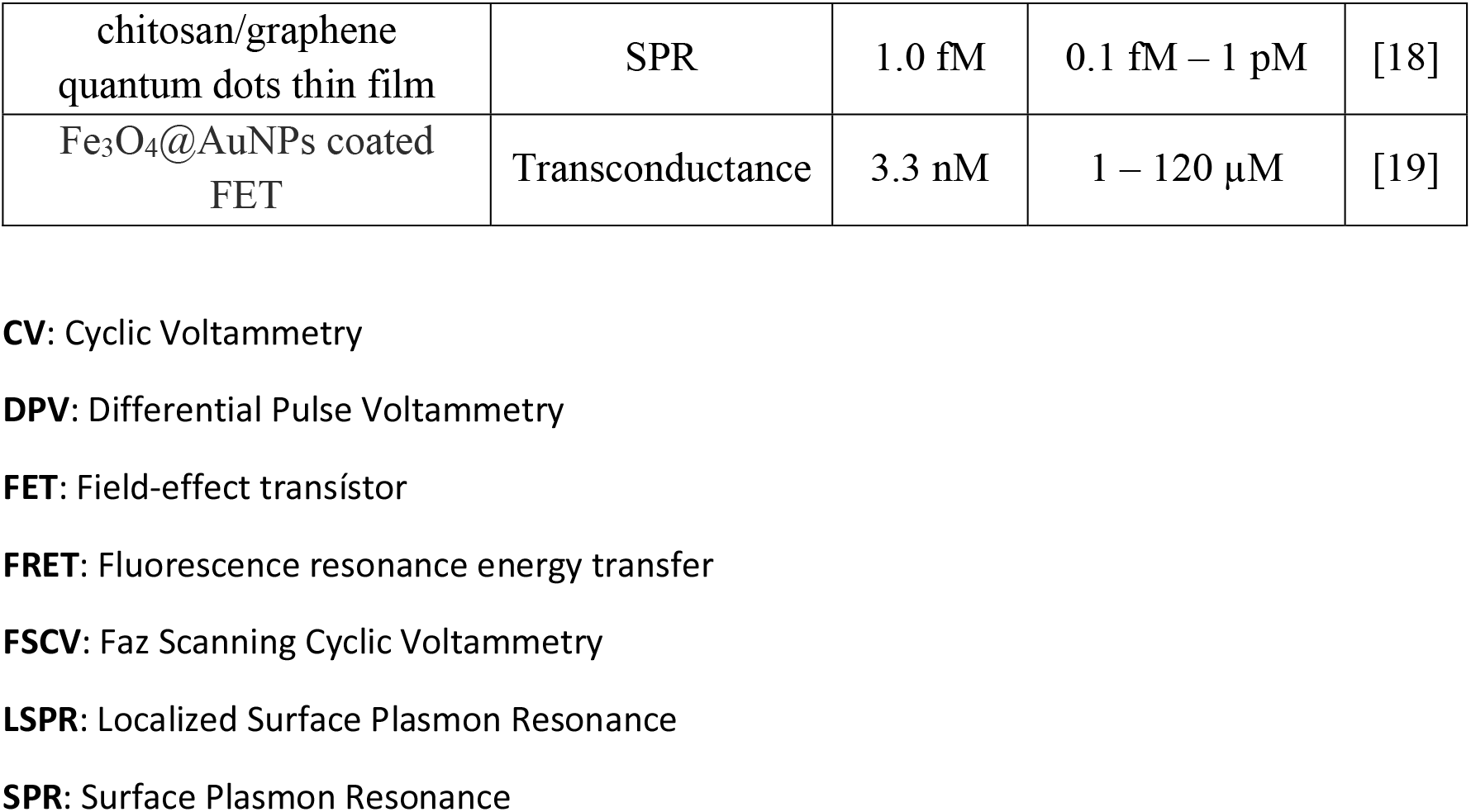
Comparison of different biosensors for dopamine detection.

**Figure S1:**
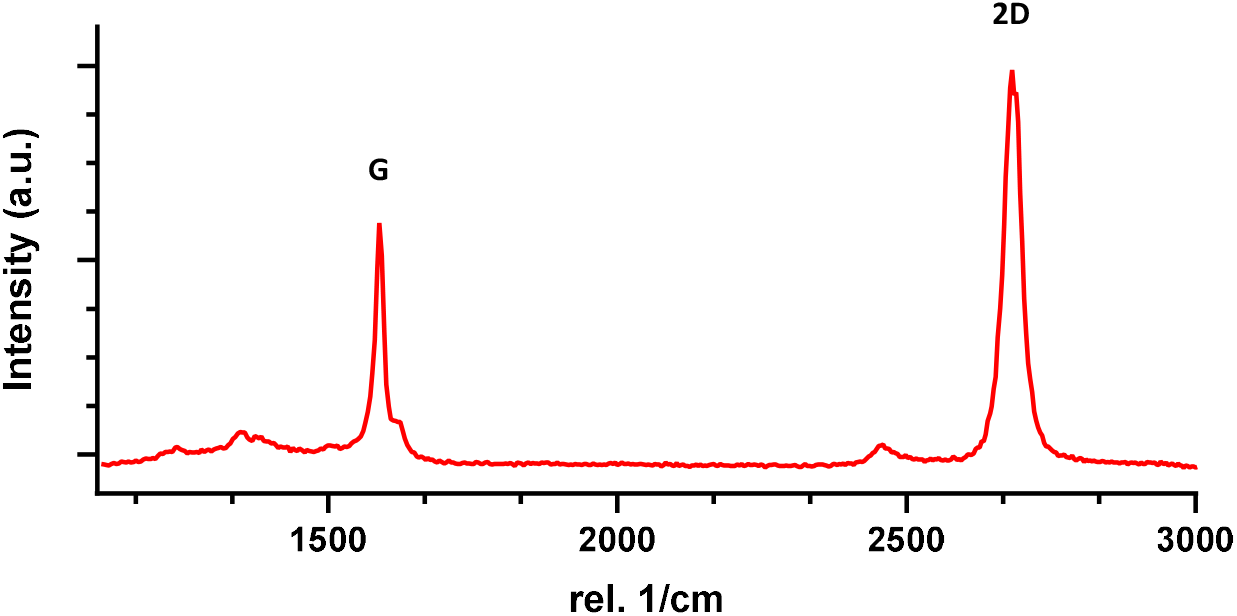
Raman spectrum of CVD grown single-layer graphene. Raman spectrum of a CVD-grown graphene sample transferred onto a Si/SiO_2_ substrate.

Raman spectroscopy was performed under laser excitation (λ = 532 nm), and spectra were acquired from 10 × 10 μm^2^ samples of CVD-grown graphene sheets transferred onto 200mm Si/SiO_2_ substrates. Figure S1 shows a characteristic Raman spectrum of a graphene sample from a graphene sheet grown for transistors’ fabrication. Two clear peaks were observed at approximately 1580 cm^−1^ and 2680 cm^−1^, corresponding to the characteristic G and 2D bands, respectively [20], [21]. A small broad peak at approx. 1350 cm^−1^, corresponding to the D band, was also observed. 2D-to-G intensity ratio was approx. 1.4. These results confirm the presence of crystalline single-layer graphene in our samples [20], [21].

**Figure S2:**
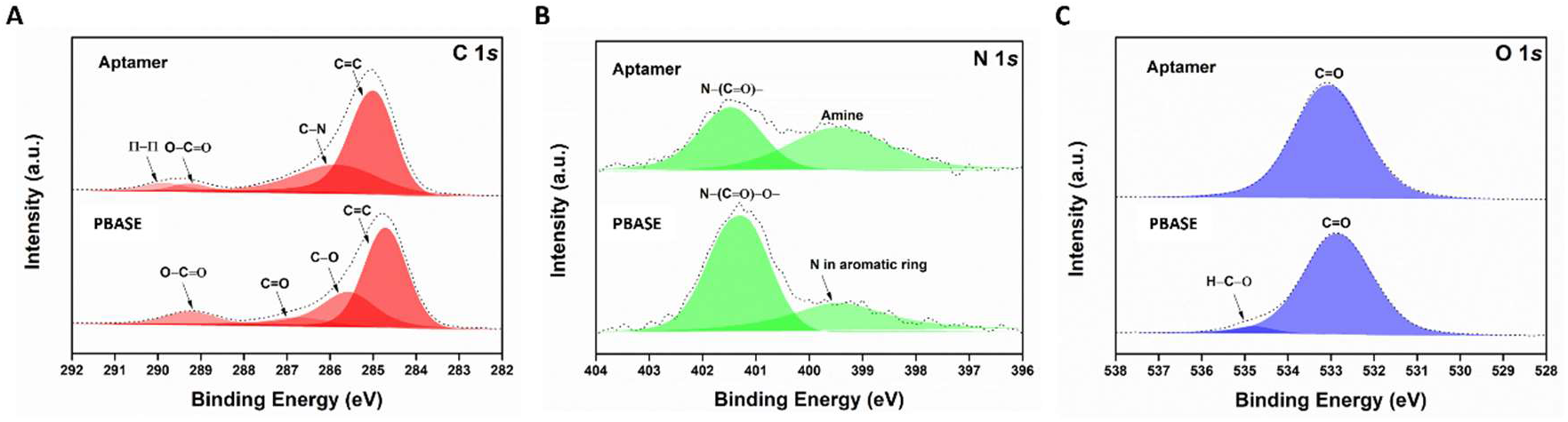
XPS peak fitting for PBASE and PBASE + aptamer samples. High-resolution XPS spectra from PBSAE and PBASE + DNA aptamer samples on graphene substrates for (A) Carbon (C 1s), (B) Nitrogen (N 1s), and (C) Oxygen (O 1s): acquired spectra (dashed lines) and peak fitting results (colored areas).

To confirm and further analyze the PBASE + aptamer bond for the implemented biofunctionalization process, high-resolution XPS spectra of C 1s, O 1s, and N 1s were fitted with Avantage data processing software (Thermo Fisher Scientific). Smart-type background subtraction was used for peak fitting, and quantification was done using sensitivity factors provided by the Avantage library. The XPS results support the successful binding of the DNA aptamer to PBASE in the substrate. Considering that the DNA aptamer binds to either the carbonyl group (C=O) or the carboxylate (C-O) part of the crosslinker [22], a reduction of these peaks, at approximately 287 eV and 285.5 eV, respectively, was observed in the PBASE + aptamer sample when compared with the PBASE sample (Fig. S2-A). The C-O bond peak from the PBASE sample became a C-N bond peak at approximately 286 eV in the PBASE + aptamer due to new contributions of C-O-C and C-OH bonds from the DNA aptamer sugar unit (Fig. S2-A) [23]. A decrease of the N-(C=O)-O-bond peak at approximately 401.5 eV and an increase of the aromatic peak at approximately 399.4 eV was observed in the PBASE + aptamer sample when compared with the PBASE sample due to the amine termination of the DNA aptamer strand (Fig. S2-B). Two peaks were observed in the O 1s spectrum, a C=O bond peak at approximately 533 eV, which is from PBASE’s aromatic ring, and an H-C-O bond peak at approximately 535 eV, which is from the PBASE’s strand that links directly to the amino group from DNA (Fig. S2-C) [23]. The H-C-O bond observed in the PBASE sample was not present in the PBASE + aptamer sample, likely due to the DNA aptamer amino group’s binding to the crosslinker. For the same reason, the C=O bond peak in the PBASE + aptamer sample increases compared with the PBASE sample (Fig. S2-A).

**Figure S3:**
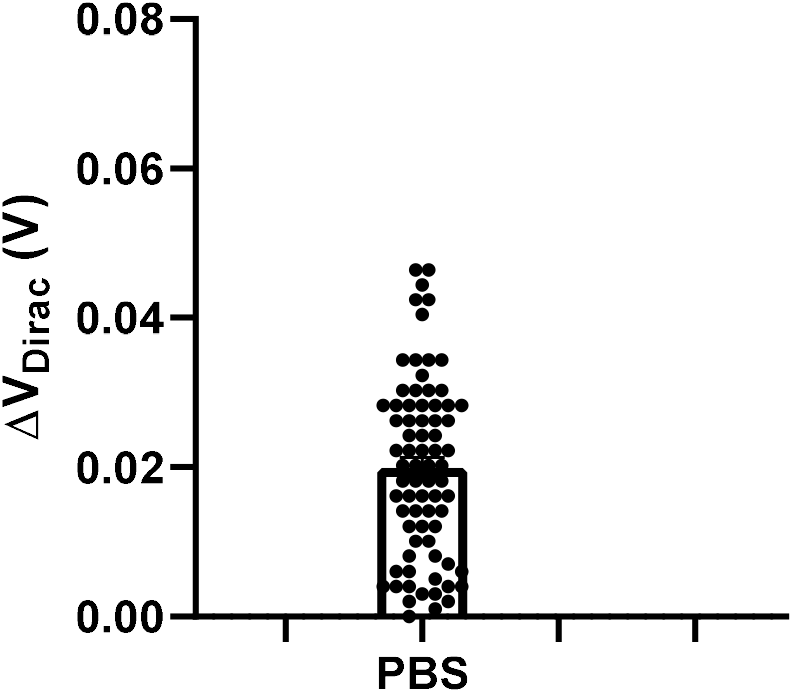
Blank samples measurements in PBS. Transconductance shifts from baseline (ΔV_DIRAC_) for blank samples not containing dopamine in 1 ×PBS (left). Data is mean ± sem.

To correctly determine the limit-of-blank of our gMTAs to offset the calibration curves, 20 μL samples not containing dopamine molecules but prepared from dopamine stock solution diluted to zM (10^−21^) in 1 × PBS, were incubated for 20 minutes. Figure S3 shows the observed transconductance shifts from baseline (ΔV_DIRAC_) for the blank samples, with an average of 19.9 ± 1.3 mV.

**Figure S4:**
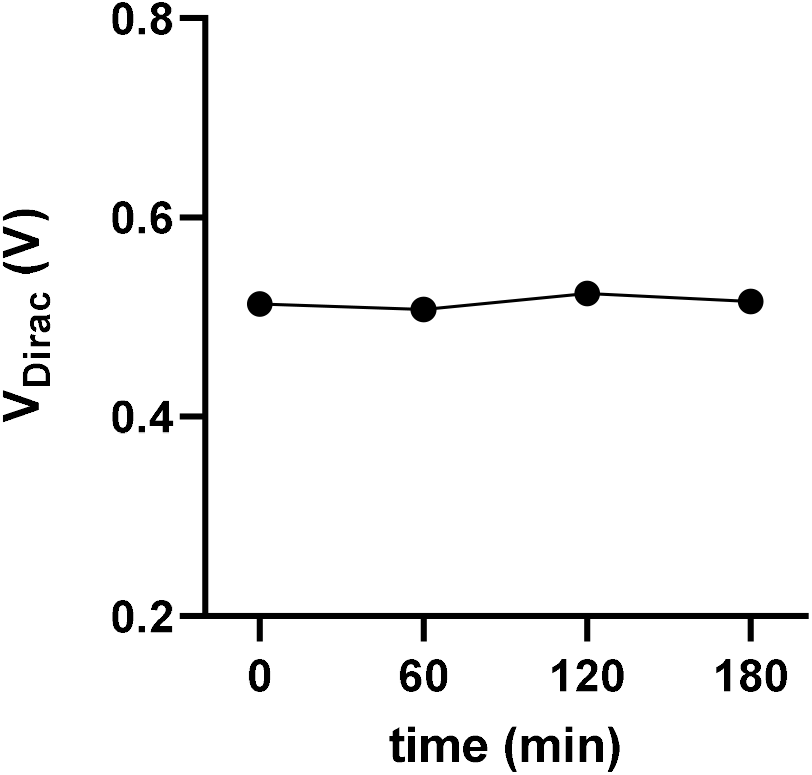
Stability of gMTA measurements. Transconductance (V_DIRAC_) measurements of PBS samples over time. Data is mean ± sem.

To assess the stability of our measurements, 20 μL PBS samples were continuously incubated in one gMTA for 3 hours, and transconductance measurements were taken every 60 min. The coefficient of variation (CV) was calculated as the standard deviation to the measurements’ mean ratio. A low CV of 1.13% was obtained, which indicates high stability.

